# Metabolic Engineering Boosts Strigolactone Production in *Nicotiana benthamiana* and Uncovers a Novel P450 Function

**DOI:** 10.64898/2026.04.02.716082

**Authors:** Meixiu Dong, Changbin Niu, Zhenrong Qiu, Xiaolong Zhong, Ralf Welsch, Ruifeng Yao, Harro J Bouwmeester, Lemeng Dong, Changsheng Li

**Affiliations:** Yuelushan Laboratory, Hunan Province Key Laboratory of Plant Functional Genomics and Developmental Regulation, Hunan Research Center of the Basic Discipline for Cell Signaling, State Key Laboratory of Chemo/Biosensing and Chemometrics, College of Biology, Hunan University, Changsha 410082, China; Plant Hormone Biology Group, Swammerdam Institute for Life Sciences, University of Amsterdam; Science Park 904, 1098 XH, Amsterdam, The Netherlands; Faculty of Biology II, University of Freiburg, 79104 Freiburg, Germany; Longping Agricultural College, Hunan University, Changsha 410082, China

## Abstract

Strigolactones (SLs) are plant hormones regulating shoot branching and symbiotic interactions, but their trace-level abundance limits research and applications. Here, we optimized a *Nicotiana benthamiana* transient expression system for SL production by tuning agroinfiltration parameters and co-expressing rate-limiting carotenoid biosynthetic genes. Overexpression of *Zea mays PSY1* or an *Arabidopsis PSY-GGPS11* fusion increased carlactone production over 2-fold and enhanced downstream SL accumulation. Using this platform, we discovered that sorghum cytochrome P450 *SbCYP728B35* catalyzes conversion of 5-deoxystrigol to sorgolactone, revealing a previously unknown function. These results establish metabolic engineering of precursor supply as an effective strategy for boosting SL production and demonstrate *N. benthamiana* as a robust system for pathway elucidation and biotechnological synthesis of bioactive strigolactones.

## Introduction

The first strigolactone (SL) strigol was identified in 1972, as a germination stimulant for *Striga lutea* (Cook et al., 1972). After a few decades, however, other functions of SLs were discovered. They promote arbuscular mycorrhizal fungi (AMF) colonization and also regulate plant branching/tillering as a “new” class of plant hormone (Akiyama et al., 2005; Gomez-Roldan et al., 2008; Umehara et al., 2008).

Thus far, more than 40 different natural SLs have been identified from plant species, consisting of canonical and non-canonical types all sharing the conserved D ring (Bouwmeester et al., 2020; Zhou et al., 2025). SLs were initially thought to be sesquiterpene lactones (Butler, 1995), however, using fluridone (carotenoid biosynthesis inhibitor), it turned out that they are carotenoid derivatives (Matusova et al., 2005). SL biosynthesis is derived from the plastidial methylerythritol 4-phosphate (MEP) pathway (Fig. 1A) (Matusova et al., 2005). In the MEP pathway, 1-deoxy-D-xylulose 5-phosphate synthase (DXS) catalyzes the condensation of D-glyceraldehyde 3-phosphate (G3P) and pyruvate, to produce 1-deoxy-D-xylulose 5-phosphate (DXP). DXP reductoisomerase (DXR) converts DXP further to MEP, which is subsequently converted to isopentenyl diphosphate (IPP) and dimethylallyl diphosphate (DMAPP), the precursors for SLs, cytokinins, sesquiterpenes and triterpenes (Vranová et al., 2013). Geranylgeranyl diphosphate synthase (GGPPS) catalyzes the conversion of these common precursors into geranylgeranyl diphosphate (GGPP), the substrate for phytoene synthase (PSY) that catalyses the rate-limiting step in carotenoid biosynthesis, from GGPP to phytoene (Burkhardt et al., 1997). Five additional enzymatic conversions result in the formation of all-*trans*-β-carotene from phytoene (Fig. 1A).

**Figure 1.**
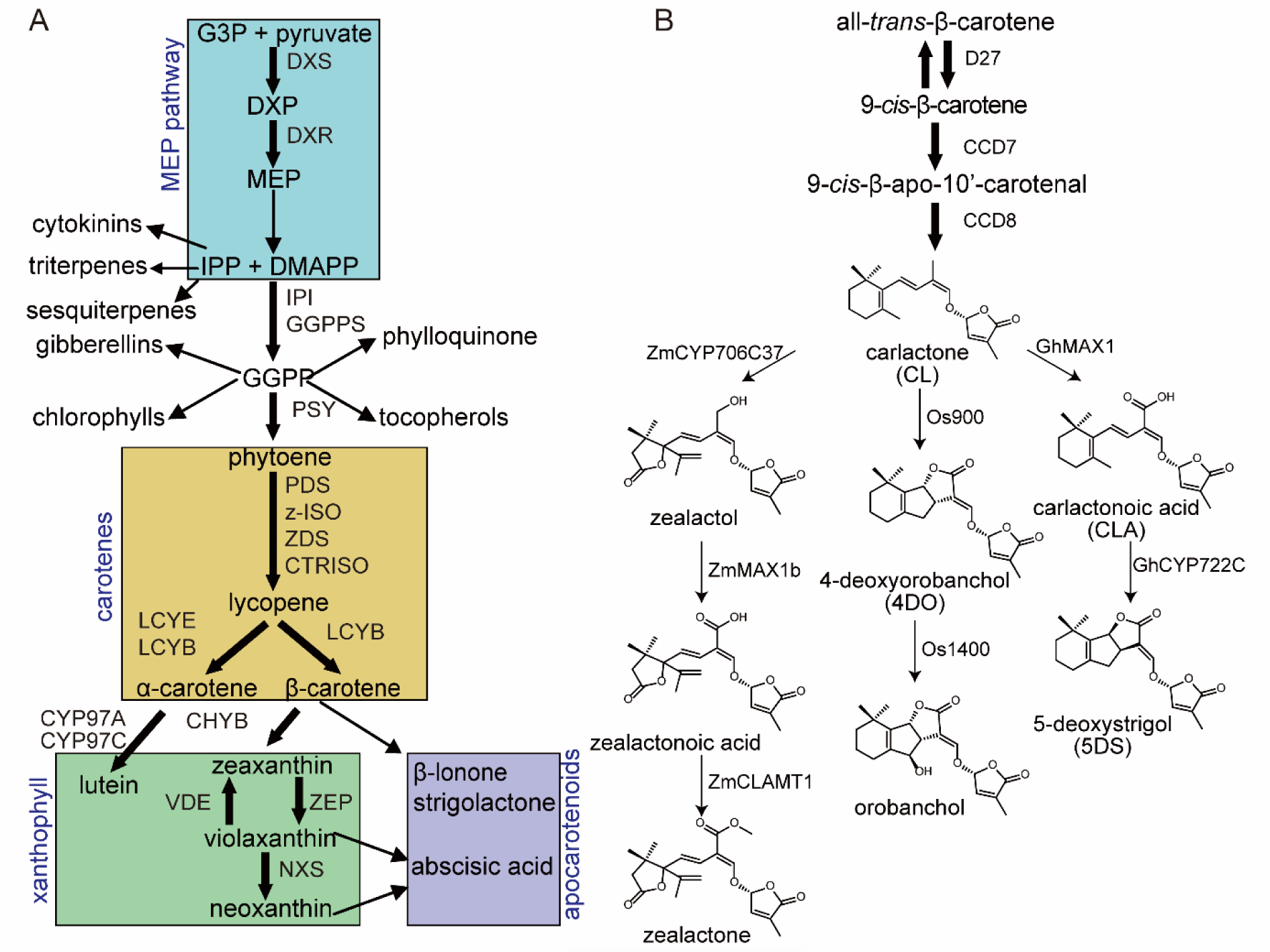
Pathways of isoprenoid biosynthesis and several strigolactones involved in this study. (A) isoprenoid biosynthetic pathways. MEP, carotenes, xanthophyll, and apocarotenoids pathways were presented in boxes with colors. G3P, glycerol triphosphate; DXP, 1-deoxy-D-xylulose-5-phosphate; MEP, 2-C-methyl-D-erythritol-4-phosphate; DXS, 1-deoxy-D-xylulose-5-phosphate synthase; DXR, 1-deoxy-D-xylulose-5-phosphate reductoisomerase; IPP, isopentenyl diphosphate; DMAPP, dimethylallyl diphosphate; GGPP, geranylgeranyl diphosphate; IPI, isopentenyl diphosphate isomerase; GGPPS, GGPP synthase; PSY, phytoene synthase; PDS, phytoene desaturase; Z-ISO, ζ-carotene isomerase; ZDS, ζ-carotene desaturase; CRTISO, carotenoid isomerase; LCYE, lycopene ε-cyclase; LCYB, lycopene β-cyclase; CHYB, β-carotene hydroxylase; CYP97C, cytochrome P450-type monooxygenase 97C; ZEP, zeaxanthin epoxidase; VDE, violaxanthin de-epoxidase; NXS, neoxanthin synthase. Names of enzymes involved are indicated along the bold arrows and the thin arrows present several steps. (B) schemes of biosynthetic pathways of carlactone (CL) and some representative downstream SLs involved in this research.

*En route* to SL biosynthesis, three enzymes are responsible for the conversion of all-*trans*-β-carotene into carlactone, the common precursor for all SLs (Alder et al., 2012) (Fig. 1B). These three enzymes *D27*, *CCD7/D17/HTD1/MAX3/RMS5/DAD3*, and *CCD8/D10/MAX4/ RMS1/DAD2* have been identified in several plant species (Morris et al., 2001; López-Ráez et al., 2008; Umehara et al., 2008; Drummond et al., 2009; Alder et al., 2012; Kohlen et al., 2012; Decker et al., 2017).

Several cytochrome P450s have been shown to be involved in the biosynthetic pathways downstream of carlactone (Niu et al.), such as MAX1 (Zhang et al., 2018). The A1 type MAX1s (MAX1 in *Arabidopsis thaliana* and its homologs from tomato/*Solanum lycopersicum* and poplar/*Populus trichocarpa*) convert carlactone into calactonoic acid, while the conversion of carlactone to 4-deoxyorobanchol (4DO) is catalyzed by A2 type MAX1s (e.g. rice/*Oryza sativa* Os900 and the Selaginella/*Selaginella moellendorffii* SmMAX1a/b) (Fig. 1B). A3 type MAX1s (rice Os1400 and maize/*zea mays* ZmMAX1b) convert carlactone to calactonoic acid and 4-DO to orobanchol (Yoneyama et al., 2018) (Fig. 1B). CYP722C is another cytochrome P450 demonstrated to be involved in SL biosynthesis. In tomato and cowpea (*Vigna unguiculata*), CYP722C catalyzes orobanchol formation from carlactonoic acid (CLA) (Wakabayashi et al., 2019), whereas the cotton (*Gossypium arboreum*) and lotus (*Lotus japonicus*) homologs (GaCYP722C, LjCYP722C) catalyze the conversion of CLA to 5-deoxystrigol (Wakabayashi et al., 2020). Moreover, we recently identified a novel cytochrome P450 in maize (ZmCYP706C37) which uses both carlactone and methyl carlactonoate (MeCLA) as substrate, to form maize SLs, zealactol and zealactone (Fig. 1B) (Li et al., 2023).

SLs exist in extremely low abundance (pmol/l of plant root exudate, fmol/g of plant root fresh weight) (Visentin et al., 2016). Additionally, the complicated structures (stereochemistry) and low stability of SLs make them difficult to be chemically synthesized. Furthermore, a great number of new SLs have been identified in the past decades (Bouwmeester et al., 2020; Yao et al., 2026), but their biosynthesis is still largely uncovered. The large biological activity of SLs makes them interesting targets, for example, as suicidal germination stimulants for parasitic plants or growth-promoting hormones for use in agriculture (Uraguchi et al., 2018). However, since the SLs are produced in very low concentrations, these activities are largely hampered. Thus, exploring suitable heterologous expression systems may contribute to addressing those issues and provide opportunities for better utilization of SLs.

*Nicotiana benthamiana* is a solanaceous species, which has been widely and increasingly used for transient expression of plant natural product biosynthetic pathways (Tian et al., 2025; Zhang et al., 2026). This system represents several advantages: agroinfiltration is relatively easy to perform, it is rapid and flexible for reconstituting different pathways or expressing various genes, and genes from a variety of plant species can be used (due to similarities in cellular compartmentalization, cofactors and coenzymes, and (partially) shared upstream precursors/substrates). Indeed, engineering strategies, such as using RNAi of competing pathways and co-expression of upstream precursor pathway genes, effectively enhanced the production of targeted isoprenoids (Rodriguez-Concepcion and Daròs, 2022). For example, the rice 4DO and orobanchol pathways were reconstituted in *Nicotiana benthamiana*(Zhang et al., 2014). However, the quantity of SLs produced by reconstitution in *N. benthamiana* is low, possibly due to low precursor availability. Effective methods to boost SL production have not been reported.

Thus, in this study we aimed to determine what are feasible methods to boost the production of SL precursor carlactone and SLs by metabolic engineering of the *Nicotiana benthamiana* transient expression factory. This could benefit the *in planta* production of structurally diversified SLs and provide the possibility for further pathway gene discovery and application of SLs in both fundamental and applied research.

## Results

### Transient production of carlactone and its derived strigolactones in *Nicotiana benthamiana*

As indicated above, all the natural SLs identified so far derive from carlactone (Alder et al., 2012). Therefore, we first attempted to produce carlactone in *N.benthamiana*. Previously, transient expression of the three carlactone pathway genes (*D27*, *CCD7*, and *CCD8*) from tomato, rice and maize resulted in the successful production of carlactone in *N. benthamiana* infiltrated leaves (Zhang et al., 2018; Li et al., 2023; Li et al., 2024). We compared the combinations of the carlactone biosynthetic genes *D27*, *CCD7*, and *CCD8* from maize and rice. LC-MS/MS analysis of leaf extracts showed that both gene variants successfully led to carlactone production in *N. benthamiana*, and no significant difference in carlactone yield was detected between the two groups (fig. S1; Fig. 2A).

**Figure 2.**
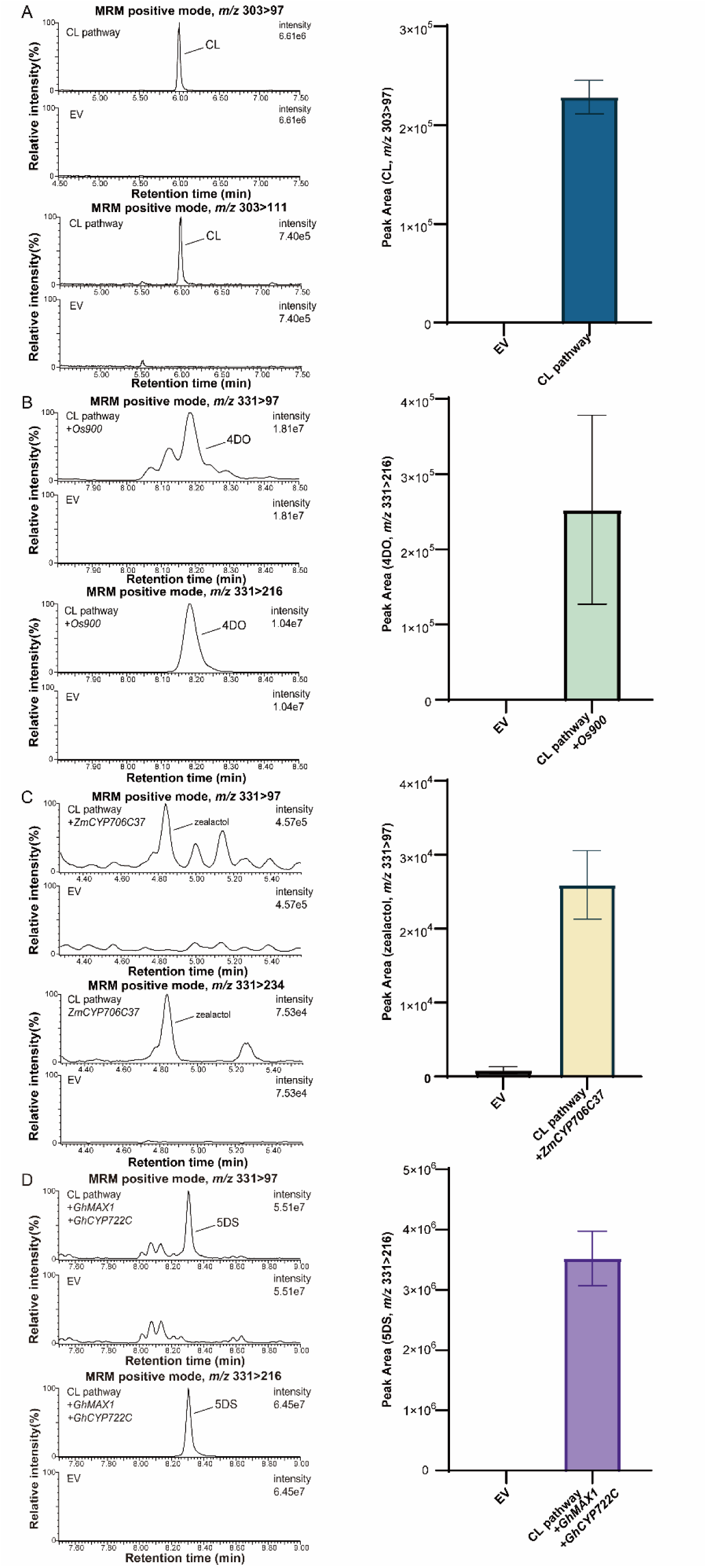
Reconstitution of carlactone and its derived strigolactones biosynthetic pathways using transient expression assays in *Nicotiana benthamiana*. (A) The detection and quantification of CL in *N. benthamiana* leaf extracts using MRM-LC–MS/MS. (B) The detection and quantification of 4DO in *N. benthamiana* leaf extracts using MRM-LC–MS/MS. (C) The detection and quantification of zealactol in *N. benthamiana* leaf extracts using MRM-LC–MS/MS. (D) The detection and quantification of 5DS in *N. benthamiana* leaf extracts using MRM-LC–MS/MS. Empty vector (EV) was used as a negative control. The data was based on at least 3 biological replicates.

Next, we attempted to use this transient plant expression system to produce other natural forms of SLs, of which the biosynthetic pathways have been elucidated. Co-infiltration of the corresponding downstream P450 genes together with the carlactone pathways resulted in production of 4DO, zealactol, and 5DS (Zhang et al., 2014; Wakabayashi et al., 2020; Li et al., 2023) (Fig. 2). These results show that this system works well with expressing several genes together and is efficient in producing natural SLs with different structures.

### Optimization of agroinfiltration methods

Previous reports have already indicated the importance of some parameters in agroinfiltration steps (Azizi-Dargahlou and Pouresmaeil, 2024). Here, we also tested some factors before and after agroinfiltration. First, three kinds of vectors were used for binary expression: one is pBI121, the other two are highly similar and modified from pBIN vector (fig. S2). The two vectors with pBIN backbone showed higher efficiency in producing SLs (4DO as an example) *in planta* than that from pBI121 (fig. S2). In next assays, we use these “pBIN” vectors. In previous projects, we noticed that the recovery status of *N. benthamiana* varies greatly after infiltration. Therefore, the effect on carlactone production of two factors, days after agroinfiltration and OD_600_ values of infiltration buffer, were observed (Fig. 3). It is clear that the amount of new-synthesized carlactone increased during three days post agroinfiltration and declined dramatically on the fourth day. But production went up again on the fifth day, although the variations among the replicated samples were relatively larger (Fig. 3A). As for the buffer concentration, the higher one (OD_600_ > 0.3) seems to be more active in inducing the production of SLs in *N. benthamiana* leaves (Fig. 3B). As we would like to search for feasible metabolic engineering strategies to boost transient heterologous production, we decided to combine “collecting sample after 3 days” and “OD_600_=0.5” in next assays, to obtain good repeatable results. It is worth noting that different labs/methods can vary a lot and these factors need to be tested and modified when designing similar projects.

**Figure 3.**
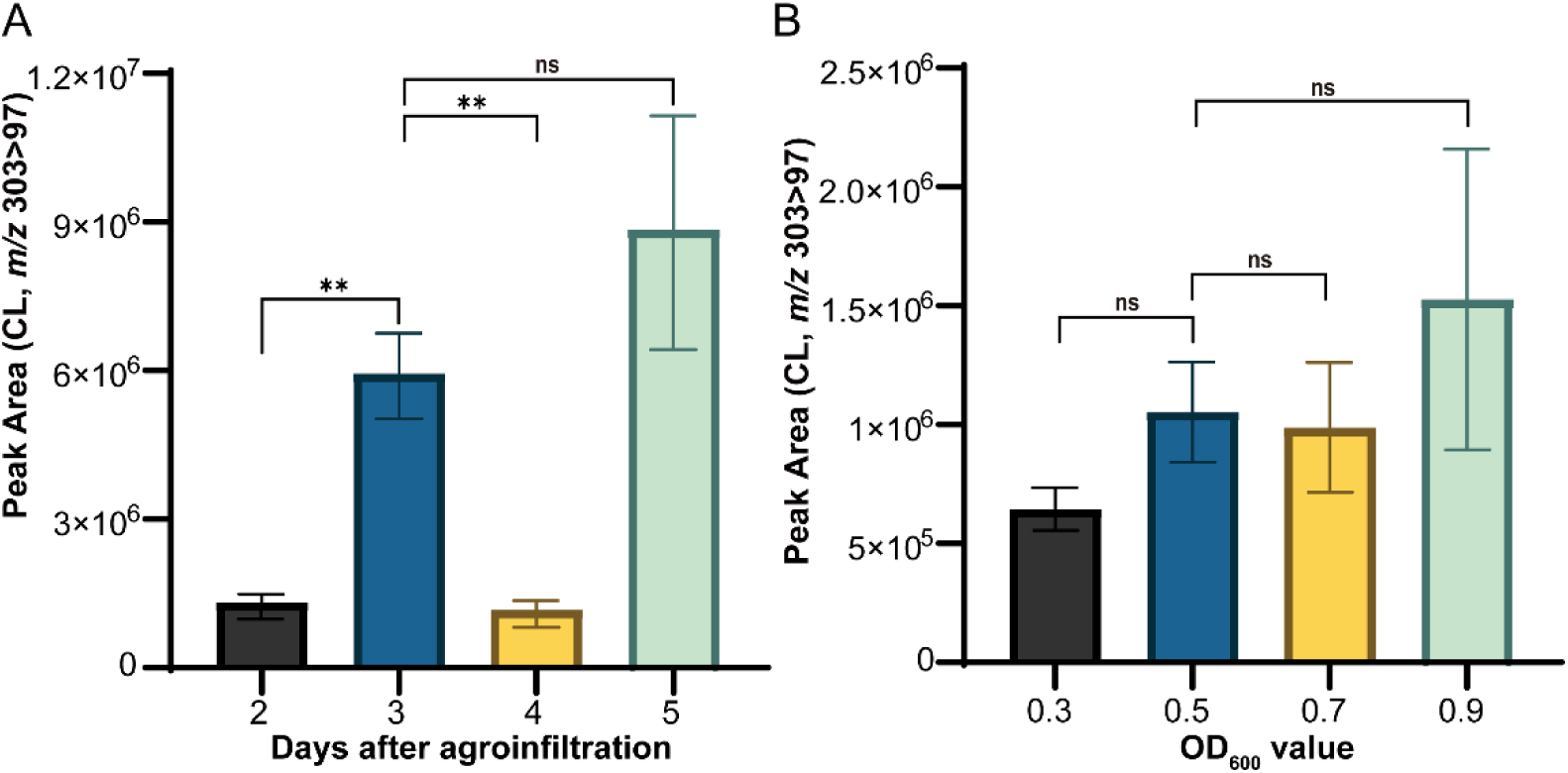
Optimization of parameters of agroinfiltration assay in the reconstitution of carlactone pathway. (A) Quantification of carlactone (CL) in *N. benthamiana* leaf extracts (peak area, transition *m/z* 303 > 97) collected from different time points after agroinfiltration. (B) Quantification of CL in *N. benthamiana* leaf extracts using different OD_600_ valuse of agroinfiltration buffer, to test the effect of concentration of agrobacterium. Empty vector (EV) was used as a negative control. The data was based on at least 3 biological replicates. Statistical analysis was performed using one-way ANOVA, where ∗ denotes significant difference (*p*<0.05), ∗∗ denotes extremely significant difference (*p*<0.01), and ns denotes no significant difference.

### Overexpression of precursor genes for β-carotene biosynthesis increases the production of carlactone

It has been shown that co-expression of upstream isoprenoid pathway genes increases the production of downstream carotenoid-derived compounds (Rodriguez-Concepcion and Daròs, 2022). We thus hypothesized that co-expressing genes involved in the upstream pathway of β-carotene together with carlactone biosynthetic genes would enhance the metabolic flux towards β-carotene and that this could increase carlactone production.

To overexpress β-carotene biosynthetic pathway genes, a number of genes from different plant species (maize, Arabidopsis, and rice) were selected (Table 1). Full length copies of *ZmDXR* (*GRMZM2G05697 / Zm00001d040163*), *ZmGGPPS1* (*AC194970.5_FG001 / Zm00001d006678*), and *ZmPSY1* (*GRMZM2G300348 / Zm00001d036345*) were cloned and co-expressed individually and in combination with the rice carlactone pathway genes. Targeted LC-MS/MS analysis was performed to analyze the carlactone amount in agroinfiltrated leave extracts. Fig. 4A shows that agroinfiltration of *ZmPSY1* alone or in combination with other genes increased carlactone production up to 2.6-fold compared with the control (expressing only rice carlactone pathway genes). Co-expressing the rice calactone pathway genes with the other two maize genes (*ZmDXR*, *ZmGGPPS1*) alone or in combination did not improve carlactone production (Fig. 4A).

**Figure 4.**
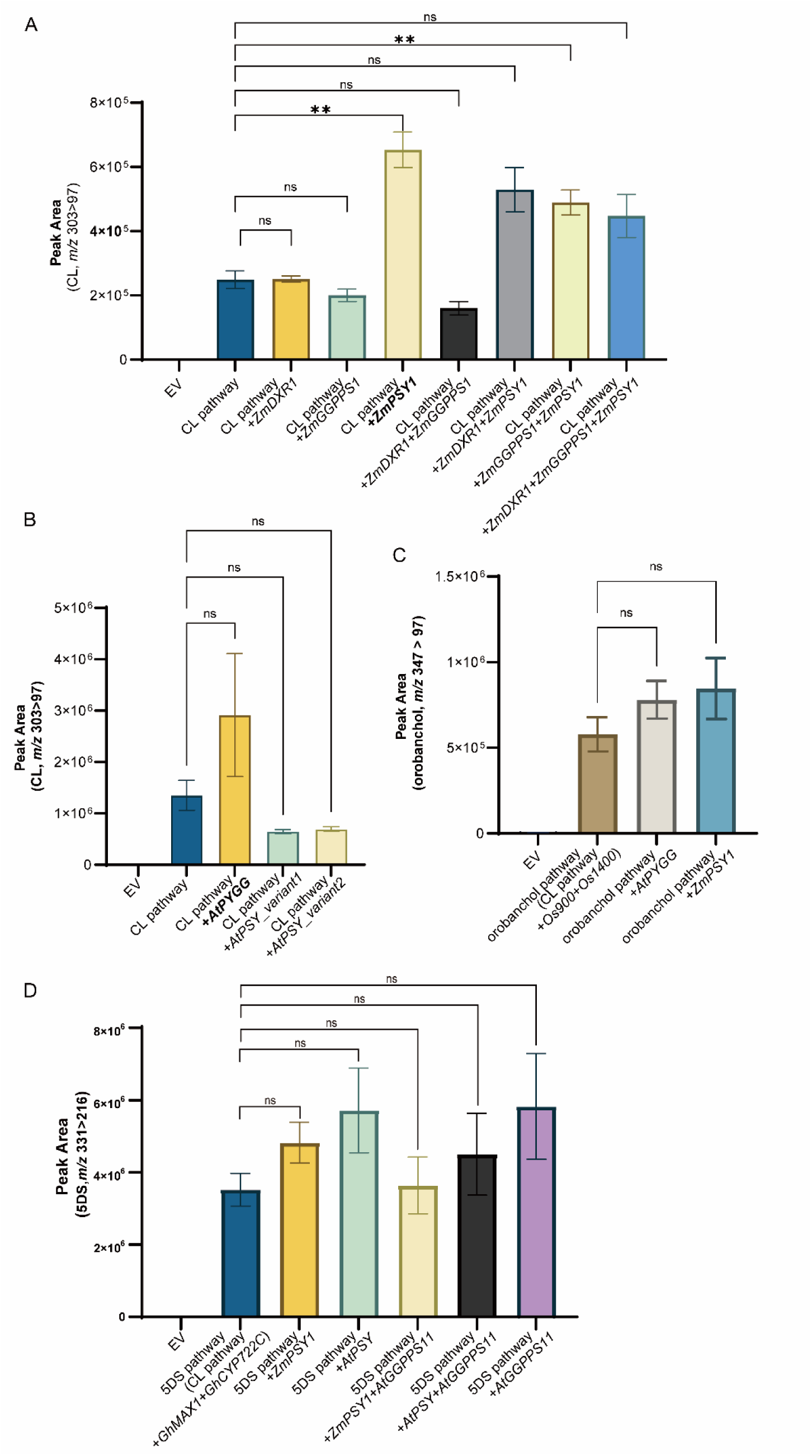
Improved production of different strigolactones in transient expression assays in *Nicotiana benthamiana* by co-expression of isoprenoid pathway genes from maize and/or *Arabidopsis*. (A) Quantification of carlactone (CL) in *N. benthamiana* leaf extracts by overexpression of isoprenoid pathway genes from maize together with CL pathway genes. (B) Quantification of carlactone (CL) in *N. benthamiana* leaf extracts by overexpression of isoprenoid pathway genes from *Arabidopsis* together with CL pathway genes. (C) Quantification of orobanchol in *N. benthamiana* leaf extracts by co-expression of effective upstream genes (selected from results in A and B panels) and orobanchol pathway genes (CL pathway genes+*Os900*+*Os1400*). (D) Quantification of 5DS in *N. benthamiana* leaf extracts by co-expression of effective upstream genes (selected from results in A and B panels) and 5DS pathway genes (CL pathway genes+*GhMAX1* +*GhCYP722C*). Empty vector (EV) was used as a negative control. The data was based on at least 3 biological replicates. Statistical analysis was performed using one-way ANOVA, where ∗ denotes significant difference (*p*<0.05), ∗∗ denotes extremely significant difference (*p*<0.01), and ns denotes no significant difference.

Also, Arabidopsis β-carotene biosynthesis pathway genes were coexpressed with the carlactone pathway genes to evaluate their effects in boosting carlactone production in *N. benthamiana*. The Arabidopsis *PSY*-*GGPS11* fusion (*AtPYGG*) increased the production of carlactone up to 2.2-fold. Interestingly, unlike the results with *ZmPSY1*, agroinfiltration of *AtPSY* together with the carlactone pathway genes did not result in an increase in carlactone production (Fig. 4B).

To confirm these results, we repeated the experiment using the two most effective genes in combination with the carlactone pathway genes and again the amount of carlactone produced doubled by co-expressing either *ZmPSY1* or *AtPYGG* with the carlactone pathway genes compared with the control, although no significant differences were found due to the variations (Fig. 4C). Furthermore, using 5DS as our target SL, we combined one or two effective genes (*ZmPSY1*, *AtGGPPS11*, *AtPSY*) with 5DS pathways genes (Wakabayashi et al., 2020) and detected 5DS from the leaf extracts. Likewise, in most cases, the addition of these upstream genes could enhance the production of 5DS, compared with the control (no significant differences) (Fig. 4D). However, the effect of combination of ZmPSY1 and AtGGPPS11 is not clear and the 5DS amount in this treatment is even lower than the samples expressing either *ZmPSY1* or *AtGGPPS11* (Fig. 4D). This suggests that the interaction or co-work of these two genes/enzymes might be a limiting factor in β-carotene and SL biosynthesis (Ezquerro et al., 2023).

### Silencing of endogenous competing pathways of β-carotene in *Nicotiana benthamiana*

Silencing of endogenous *N. benthamiana* genes involved in competing pathways is another strategy to increase target products, and this has previously been reported to improve (+)-valencene production (Cankar et al., 2015). To further direct the metabolic flux through the β-carotene pathway, we silenced the expression of two genes (*NbLCYE*, *NbCHYB*), which encode enzymes that compete for lycopene and β-carotene, respectively. Fragments of *NbLCYE* and *NbCHYB*1 were cloned into the RNAi silencing vector. However, simultaneous overexpression of the rice carlactone pathway genes and RNAi silencing of *NbLCYE* and/or *NbCHYB1* did not contribute to any enhancement of carlactone production (fig. S3).

### Discovery of new function of *SbCYP728B35* in sorghum strigolactone biosynthesis

In this project, we also combined *SbCYP728B35* together with 5DS pathway and expect to produce sorgomol in *N. benthamiana*, of which the function was revealed (Wakabayashi et al., 2021). However, no clear peak/signal of sorgomol was detected from this sample and a new compound with *m/z* 317 in ESI positive mode was highly accumulated. From published data, this compound could be sorgolactone (Hauck et al., 1992; Sugimoto et al., 1997). To validate this, we grew Jingxi, a *Sorghum bicolor cv. Dochna* commercial variety and collected its root exudate after low-phosphorus treatment to enhance the SL production. It was shown that this sorghum line produced high amount of 5DS and similar compound as found in *N. benthamiana* sample expressing 5DS pathway and *SbCYP728B35*. Daughter ion scanning of this compound from these two types of samples showed that they are identical in structures (Fig. 5, A and B), which are highly possible to be sorgolactone or similar strigolactons.

**Figure 5.**
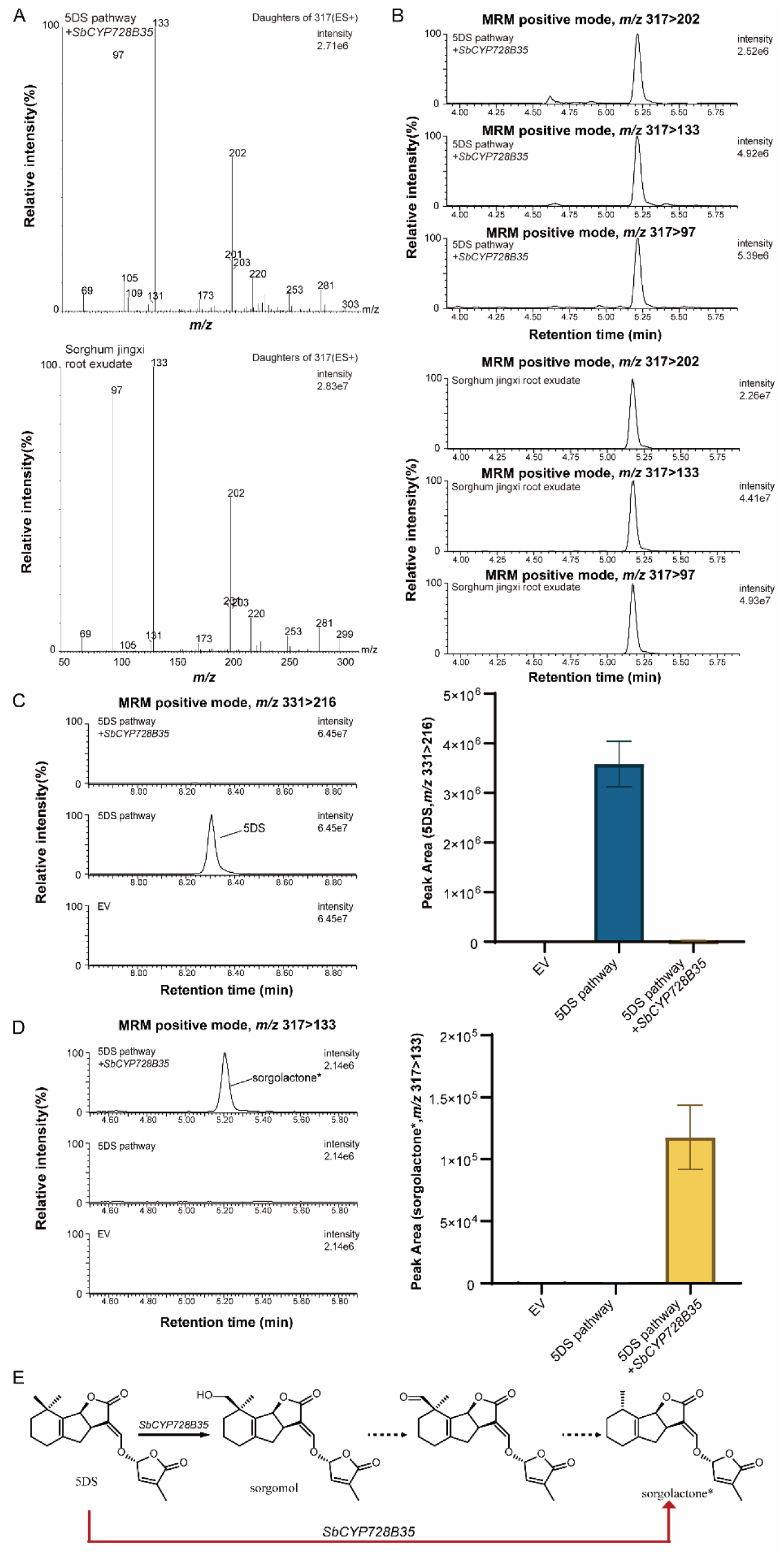
Identification of new function of a P450 gene in sorghum strigolactone biosynthesis. (A) MS-MS fragmentation spectra of natural putative sorgolactone from sorghum root exudate and *N. benthamiana* leaf extracts expressing 5DS pathway and a sorghum gene *SbCYP728B35*. (B) MRM methods/channels for detection of this putative sorgolactone using LC-MS/MS. (C) Quantification of 5DS in *N. benthamiana* leaf extracts over-expressing 5DS pathway genes (CL pathway genes+*GhMAX1* +*GhCYP722C*) with or without co-expression of *SbCYP728B35*. (D) Quantification of this putative sorgolactone in *N. benthamiana* leaf extracts over-expressing 5DS pathway genes (CL pathway genes+*GhMAX1* +*GhCYP722C*) with or without co-expression of *SbCYP728B35*. (E) Scheme for function of *SbCYP728B35* in strigolactone biosynthesis. The black solid arrow indicates the previously published activity of SbCYP728B35 while the black dash arrows represent our assumed steps. The red arrow indicates the new function of *SbCYP728B35* in converting 5DS to putative sorgolactone based on our result. Empty vector (EV) was used as a negative control. The data was based on at least 3 biological replicates.

Moreover, 5DS and this putative sorgolactone were also detected and quantified from different *N. benthamiana* samples (Fig. 5, C and D). It is obvious that 5DS was consumed in the sample expressing *SbCYP728B35* and no putative sorgolactone was found in 5DS-producing leaf extract. This led to our hypothesis of sorgolactone biosynthesis, which is catalyzed by SbCYP728B35 (Fig. 5E). In previous study, although the enzymatic activity of this enzyme in catalyzing the conversion of 5DS to sorgomol was characterized, the detection or biosynthesis of this sorgolactone has not been reported in recent years. We presented that SbCYP728B35 itself could catalyze this putative sorgolactone production from 5DS, but the involvement of sorgomol in sorgolactone biosynthesis can still be explored (Fig. 5E).

## Discussion

SLs, the most recently discovered plant hormone, play significant roles in plant development and the interaction of plants with other organisms. As a plant hormone, SLs interplay with other phytohormones, to modulate growth and development. In the root rhizosphere, SLs act as signaling molecules in the communication of plants with beneficial as well as detrimental organisms (López-Ráez et al., 2017). However, their extremely low concentration in natural conditions hinders the discovery of full biosynthetic pathways and potential applications (such as combating parasitic plants such as *Striga*/witchweed through suicidal germination).

In the present study, we showed that it is possible to optimize the expression platform in *N. benthamiana* to produce SLs and their precursor carlactone. Here, three approaches were undertaken, modification of the agroinfiltration method, heterologous overexpression of β-carotene pathway genes and silencing of competing pathways. The first one gave us hint about the sample collection time points, the vector, and the effect of buffer (Fig. 3 and fig. S2). Among the tested candidates in the second strategy, the overexpression of either *AtPYGG* or *ZmPSY1* together with the rice carlactone genes resulted in more than 2-fold higher accumulation of carlactone (Fig. 4), suggesting that these two genes indeed promote the upstream metabolic flux through the precursor pathway. *ZmPSY1* was used in the ‘Golden Rice 2’ project, in which a higher β-carotene accumulation was also achieved by this gene (Paine et al., 2005). The roles of these two genes (*AtPYGG* and *ZmPSY*) in boosting SL production were further confirmed by combining them with the entire orobanchol and 5DS pathway (Fig. 4). In this way, around 2-and 3-fold higher levels of orobanchol and zealactone were achieved, respectively. This proves that co-overexpression of a complicated pathway involving multiple genes is feasible in *N. benthamiana* transient expression. Since β-carotene is the precursor for carlactone and all the other SLs, we speculated that the amount of β-carotene might be a bottleneck for further increments. In previous studies, the most representative example of increasing its content was shown in ‘Golden Rice 2’, where a 17-fold increase was observed in rice endosperm overexpressing *ZmPSY1* (Paine et al., 2005). Moreover, overexpression of upstream genes (*DXS*, *PSY/* bacterial *phytoene synthase crtB*, or *lycopene β-cyclase*/*LCYb*) by transgenic engineering also resulted in (higher) accumulation of β-carotene in soybean, banana, potato tubers, and tomato fruits (Ducreux et al., 2005; Schmidt et al., 2015; Paul et al., 2017; Diretto et al., 2020). All these results suggest that overexpression of rate-limiting steps upstream in the precursor pathway is an effective strategy for enhancing carotenoid content in various plant tissues and species.

Transcript levels of *ZmDXR*, *ZmGGPPS1*, and *ZmPSY1* were found to be positively correlated with the carotenoid content in maize seeds (Vallabhaneni and Wurtzel, 2009), and experiences already proved that there is a physical interaction between PSY and GGPPS(Rao et al., 2024). So we expected that coexpression of one or more of these three genes would increase carlactone production. However, only co-expression of *ZmPSY1* resulted in an obvious increment in carlactone production, while the combination with *ZmGGPPS1* did not further raise carlactone production (Fig. 4A). This might be caused by sub-optimal interaction between ZmGGPPS1 and ZmPSY, which does not increase GGPP supply to the carlactone pathway. In plants, several copies of *GGPPS* have evolved and their activities differ, and only specific GGPPSs were shown to interact with PSY. For example, 12 and 7 *GGPPS* members were identified in Arabidopsis and *N. tabacum*, respectively, and only one of them (AtGGPPS11 and NtGGPPS1) is responsible for channeling GGPP into the production of carotenoids by interacting with PSY (Ruiz-Sola et al., 2016; Dong et al., 2022). It is thus possible that we need a different *GGPPS* (possibly from a dicot species) and/or *PSY* to better channel GGPP to carlactone production in the *N. benthamiana* expression system. Indeed, upon coexpression of the Arabidopsis *PSY-GGPS11* fusion construct we do see an increase in carlactone production. This is likely due to *AtGGPPS11* alone or to the better interaction with AtPSY, as coexpression of *AtPSY* alone with the carlactone pathway genes does not increase carlactone production (Fig. 4B).

Inhibition of competing endogenous pathways has been shown to be a feasible approach in terpenoid engineering of *N. benthamiana*. For instance, by simultaneously silencing *N. benthamiana* squalene synthase (SQS) and sesquiterpene (5-*epi*-aristolochene) synthase, the transient heterologous production of the sesquiterpene, (+)-valencene was increased (Cankar et al., 2015). In another example, it was demonstrated that taxadiene production was enhanced via silencing of the *N. benthamiana PSY*, which competes with taxadiene synthase for the common substrate GGPP (Hasan et al., 2014). Here, to increase the flux into the carlactone pathway, we chose *NbLCYE* and *NbCHYB* genes as targets for silencing, which should eliminate the divergence from lycopene to α-carotene and therefore increase β-carotene production (fig. S3). Knockdown of *LCYE* or *CHYB* expression has been shown to increase total carotenoid and β-carotene contents in several plant species (Pogson and Rissler, 2000). However, in this study, silencing of these two targets (*NbLCYE*, *NbCHYB1*) did not improve carlactone production (fig. S3). Silencing of *NbLCYE* and *NbCHYB1* resulted in a significantly lower level of carlactone compared with the non-silencing control (fig. S3). One explanation could be that other competing/branching pathways downstream of lycopene or β-carotene exist, which consume the higher accumulated substrate and thus negate the effect of silencing. More detailed experiments (RT-qPCR, metabolomics, etc.) are needed to investigate the different possibilities, which would help to reveal the mechanism and improve these techniques.

Other heterologous expression systems were also used for the production of SLs or for other plant natural products (Tian et al., 2025; Zhang et al., 2026). Recently, a microbial platform to produce SLs using a bacterium-yeast consortium was developed (Wu et al., 2021). In Wu’s study, biosynthesis of SL precursor carlactone was established in *E. coli* and downstream P450s were expressed in *S. cerevisiae*. The former technique in *E. coli* was first used by Alder et al., (2012) who discovered the carlactone biosynthesis pathway. Three canonical SLs were produced in the *E. coli*–yeast consortia, showing its potential for producing SLs and characterizing P450 functions. The microbial expression system is relatively easy and clean for later isolation and purification of the compounds produced, while *N. benthamiana* is probably more flexible for overexpression of plant genes. Specifically, because of similarities in cellular compartmentalization, cofactors and coenzymes, and similar upstream precursors, little or no modifications of targeted gene sequences are required compared with bacterial or yeast systems. Moreover, multiple genes/steps in the same pathway can be easily reconstituted in *N. benthamiana*.

Several other strategies for engineering in this transient expression system might also be efficient to further boost the SL production, such as the utilization of transcription factors regulating the expression of isoprenoid pathways and exploitation of genome editing (Rodriguez-Concepcion and Daròs, 2022). Nevertheless, not all these approaches would be generally applicable in the production of various products. With more knowledge and techniques being developed currently and in the future, this transient expression *in planta* could be a promising technology, to produce SLs and also other compounds with bioactivities and commercial and medical values (Tian et al., 2025; Zhang et al., 2026).

In conclusion, we developed an optimized *Nicotiana benthamiana* platform for strigolactone production through agroinfiltration parameter tuning and metabolic engineering of precursor supply. Co-expression of *ZmPSY1* or *AtPYGG* with carlactone pathway genes boosted production over 2-fold, establishing precursor availability as a key bottleneck. This enhanced system enabled functional discovery of *SbCYP728B35* in sorgolactone biosynthesis, demonstrating its utility for pathway elucidation. Our work provides a scalable strategy for in planta synthesis of bioactive strigolactones.

## Materials and Methods

### Plant Material

*Nicotiana benthamiana* plants were grown in square pots in the greenhouse at 20-22°C with 65% humidity. After 4-5 weeks (around four to six leaves stage), plants were used for agroinfiltration.

Maize (inbred line B73) seeds were germinated on wet filter paper and the seedings were transferred to pots filled with river sand. The maize plants were grown in a compartment with 12 h 30°C:12 h 28°C photoperiod. Artificial light of 600 W/m2 was used to supplement low natural light conditions. After watering with half-strength Hoagland solution for one week, phosphate starvation was applied by removal of K_2_HPO4 (López-Ráez et al., 2008; Mohemed et al., 2016). Similar methods were applied to *Oryza sativa* ssp. *Japonica* Nipponbare, and *Arabidopsis thaliana* ecotype Col-0. Root and leaf tissues were collected for RNA extraction and cDNA synthesis.

Seeds of *Sorghum bicolor cv. Dochna* termed Jingxi was purchased from Zhengzhou Huafeng Grass Industry Technology Co., Ltd. After treatment with low phosphate, the root exudate was collected for LC-MS/MS analysis.

### Cloning and Transformation

Total RNA from leaves from plants (*N. benthamiana,* maize, rice, and *Arabidopsis*) was extracted using the TRIZOL method. Around 100 mg fresh tissue was ground into a fine powder under liquid nitrogen and 1 mL Trizol was added. After 5 min incubation, 200 µL chloroform was added and the tubes were vortexed and then inculcated for 3 min. The upper aqueous phase was obtained after centrifuging and transferred to a new tube. Isopropanol (500 µL) was used to precipitate the RNA. After washing with 1 mL 70% ethanol, the RNA pellet was air-dried and dissolved in 50 µL Rnase-free water. Removal of DNA was carried out using TURBO DNA-*free*™ Kit (Invitrogen). Afterwards, RevertAid First Strand cDNA Synthesis Kit (Thermo Scientific) was used for cDNA synthesis. The maize carotenoid genes (*ZmDXR*, *ZmGGPPS1*, *ZmPSY1*) were amplified from maize cDNA. Four plasmids containing Arabidopsis or rice/*Oryza sativa* carotenoid pathway genes were provided by Ralf Welsch. Primers with specific restriction enzyme digestion sites or homologous recombination sites were designed to re-clone coding regions of our targeted genes from cDNA or plasmids. Some of these PCR products were purified, enzyme digested and then ligated into the pIV2.1 vector. After confirmation by sequencing, LR reactions were performed to get these fragments into the binary expression vectors (Zhang et al., 2014). Some of these PCR products were purified and ligated into the binary expression vectors using in-fusion cloning method.

The rice genes/constructs *OsD27*, *OsCCD7*, *OsCCD8*, *Os900*, and *Os1400* were described before (Zhang et al., 2014). The maize carlactone genes were same as previous study (Li et al., 2023).

*N. benthamiana* genes, *NbLCYE* (Niben101Scf18343g00013.1), *NbCHYB1* (Niben101Scf01232g03010.1) and *NbCHYB2* (Niben101Scf02285g01028.1) were identified in the *Nicotiana benthamiana* draft genome sequence v1.0.1 (Sol Genomics Network) by BLAST using the *Nicotiana tabacum* genes (XM_016611548.1, NM_001325834.1, NM_001326092.1) as query (Bombarely et al., 2012). For gene silencing, the SGN VIGS Tool was used to find the sequence region for designing primers and fragment (200-250 bp) cloning (Van Bel et al., 2022). Two-step Gateway cloning was applied to create entry clones (pDONR207 containing the gene fragment) that were then subcloned into the pK7GWIWG2(II) vector (Karimi et al., 2002; Cankar et al., 2015).

Subsequently, constructs confirmed by sequencing were transformed into *Agrobacterium tumefaciens* strain AGL0 or AGL1 by heat shock.

### Transient Expression by agroinfiltration

*N. benthamiana* plants of around 4-weeks old were used for transient expression. *Agrobacterium* cultures were prepared and collected by centrifuge. The pellets were redissolved in infiltration buffer (50 mM MES-KOH with pH 5.7, 0.5 % glucose, 2 mM NaH_2_PO_4_, 200 mM acetosyringone) to adjust the OD_600_. *Agrobacterium* strains harboring the desired expression vectors were combined, and a strain transformed with the empty vector was included to equalize the concentration of strains in different combinations. In most of the experiments, AGL0 harboring *P19* was used for better overexpression by suppressing gene silencing except when we investigated transient RNAi silencing. The infiltration process has been described before. A total of three or five plants were used as biological replicates. The infiltrated leaves were collected 6 days after infiltration, snap-frozen in liquid nitrogen and stored at −80 °C until further sample preparation.

### Sample preparation and LC-MS/MS analysis

For SL extraction from *N. benthamiana* leaves, samples were ground to a powder in liquid nitrogen in a mortar and pestle. 200 mg of this powder was weighed out and transferred to a 2 mL Eppendorf vial to which 1.5 mL ethyl acetate was added. After vortexing, sonication, and centrifugation, the supernatant was transferred into a 4 mL glass vial. The pellet was extracted once more, and the supernatants were combined and dried in a SpeedVac. After drying, the residue was dissolved in 37.5 μL of ethyl acetate and 3 mL hexane. Then 2 mL ethyl acetate and 4 mL hexane were loaded onto a Strata® SI-1 Silica column (55 µm, 70 Å), 200 mg/3 mL) for equilibration. After adding the sample dissolved in 37.5 μL of ethyl acetate and 3 mL hexane, the column was washed with 2 mL of hexane and the SLs eluted using 2 mL hexane/ethyl acetate (1:9). The solvent was evaporated by vacuum and the residue dissolved in 200 μL acetonitrile/water (1:3) and filtered with a polypropylene micro-centrifugal filter (Thermo Scientific™) before analysis on LC-MS.

For SL analysis, the Acquity UPLC System (Waters, Milford, MA, USA) was utilized, which was equipped with a Xevo TQ-XS triple-quadrupole mass spectrometer (Waters MS Technologies) with electrospray ionization (ESI) interface as described before (Floková et al., 2020), with small modifications. 5 μL sample was injected onto the Acquity UPLC™ BEH C18 column (2.1 × 100 mm, 1.7 μm particle size, Waters, Milford, MA, USA). The carlactone standard was kindly provided by Professor Salim Al-Babili (King Abdullah University of Science and Technology). For CL detection, the column temperature was 45 °C and the flow rate was 0.4 mL/min. The gradient for 15 mM formic acid in water (A) and 15 mM formic acid in acetonitrile (B) was set as follows: elution for 0.5 min with 5% B, increase to 60% B at 2 min, followed by an increase of B to 90% at 7.3 min, and a return to starting condition (5% B) in 1.2 min and 1.5 min for equilibration. For SL detection (orobanchol, zealactone), the column temperature was 45 °C and the flow rate was 0.45 mL/min. The elution profile for A and B was set as follows: isocratic elution for 0.4 min with 15% B, followed by a linear increase to 27, 40 and 65% of B at 0.65, 5 and 8 min, respectively. After another 0.7 min, the percentage of B increased to 95% at 9.5 min, which was maintained for 0.8 min. At 10.5 min, the gradient was changed to 85% A/15% B, followed by a 1.5 min equilibration. The eluate was introduced in the ESI ion source of the MS under the following conditions: source/desolvation temperature (120/550 °C), cone gas flow (150 L/h), desolvation gas flow (1000 L/h), collision gas flow (0.15 ml/min), capillary/cone voltage (1.2 kV/15-20 V), collision energy (10-25 eV). MS data were recorded in multiple reaction monitoring mode (MRM) mode (Cardoso et al., 2014; Zhang et al., 2014; Charnikhova et al., 2017, 2018). MassLynx software (V4.2, Waters, Milford, MA, USA) was used to run the system and analyze the data.

## Conflict of interest statement

All the authors declare that they have no conflict of interest.

## Acknowledgments

We would like to thank our lab members for providing help in plant growing, sample preparation and other technical supports.

## Funding

This work was supported by Yuelushan Laboratory Breeding Program (YLS-2025-ZY04002, YLS-2025-ZY03001, YLS-2025-ZY01004), National Natural Science Foundation of China (32400258), Outstanding Youth Project of the Natural Science Foundation of Hunan Province (2024JJ2013), National Key Research and Development Program of China (2023YFD1401100), Hunan Science and Technology Innovation Plan (2025ZYJ003), the European Research Council (ERC) Advanced grant CHEMCOMRHIZO (670211), the Dutch Research Council (NWO/OCW) Gravitation programme Harnessing the second genome of plants (MiCRop) (024.004.014), Foreign Expert Project “High-level Foreign Expert Introduction Plan” by Ministry of Science and Technology of China (Z202304492179), and Hunan Provincial Innovation Foundation For Postgraduate (CX20240386).

## Author contributions

L.D, H.J.B and C.L. designed this study. M.D, C.N, Z.Q. and C.L. performed the experiments, analyzed the data and wrote the manuscript. X.Z. assisted in some experiments. R.W. constructed some plasmids. R.W, R.Y, H.J.B, L.D, and C.L. discussed this project. L.D, H.J.B and C.L. revised the manuscript. All authors have read and approved this manuscript.

## Notes

### Competing Interest Statement

The authors have declared no competing interest.

