## Supplemental figures for "Metabolic Engineering Boosts Strigolactone Production in *Nicotiana benthamiana* and Uncovers a Novel P450 Function"

### Supplementary files

Table 1. Targeted genes in isoprenoid biosynthetic pathways for overexpression or silencing

| Gene name/accession | Note | Reference |
|---|---|---|
| *ZmDXR* /*GRMZM2G056975* /*Zm00001d040163* | transcript level positively correlated with endosperm carotenoid accumulation; high expression level in leaves and mycorrhizal roots; the first committed step toward IPP and DMAPP formation | (Hans et al., 2004; Vallabhaneni and Wurtzel, 2009) |
| *ZmGGPPS1* /*AC194970.5_FG001* /*Zm00001d006678* | transcript level positively correlated with endosperm carotenoid accumulation; highest expression among the four maize GGPPS in leaves | (Vallabhaneni and Wurtzel, 2009) MaizeGDB expression data |
| *ZmPSY1* /*GRMZM2G300348* /*Zm00001d036345* | transcript level positively correlated with endosperm carotenoid accumulation; substantially increased carotenoid accumulation (preferential β-carotene) in "Golden Rice 2" | (Paine et al., 2005; Vallabhaneni and Wurtzel, 2009) |
| *AtPYGG* (Arabidopsis *PSY-GGPS11* metabolon, from construct *pRSF-PSY-GGPS*) | the fusion protein is stable in plants and not subjected to nonspecific or targeted protein degradation; the fusion is capable of increasing the IPP conversion efficiency specifically toward the synthesis of carotenoids | (Camagna et al., 2019) |
| *AtPSY* (from construct *pRSF-AtPSY*) | A187D exchange in Arabidopsis PSY resulted in higher enzymatic activity (higher lycopene production) than the wild type version | (Welsch et al., 2010) |
| *AtPSY_variant* (from construct *pRSF-AtPSY_AtoD*) | | |
| *NbLCYE* / *Niben101Scf18343g00013.1* | to silence the competing pathways of β-carotene | This study |
| *NbCHYB1* / *Niben101Scf01232g03010.1* | to silence the competing pathways of carlactone | This study |
| *NbCHYB2* / *Niben101Scf02285g01028.1* | to silence the competing pathways of carlactone | This study |

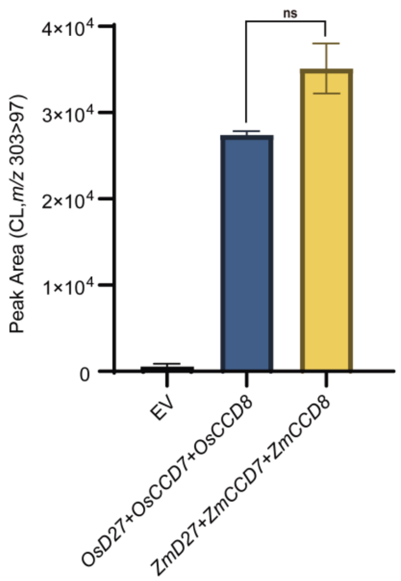


**figure S1. Comparation of carlactone production in *Nicotiana benthamiana* when expressing rice or maize carlactone pathway genes.** Quantification of carlactone (CL) in *N. benthamiana* leaf extracts (peak area, transition *m/z* 303 > 97) transiently produced by co-infiltration of rice or maize CL pathway genes. Empty vector (EV) was used as a negative control. The data was based on at least 3 biological replicates. Statistical analysis was performed using one-way ANOVA, where ∗ denotes significant difference ($p$<0.05), ∗∗ denotes extremely significant difference ($p$<0.01), and ns denotes no significant difference.

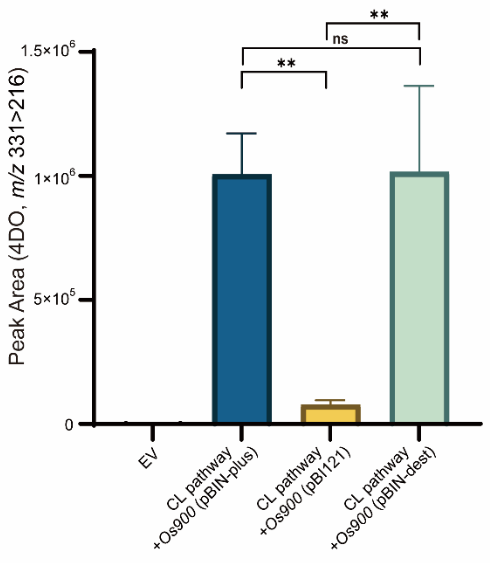


**figure S2. Comparation of 4DO production in *Nicotiana benthamiana* when using different binary expression vectors.** Quantification of 4DO in *N. benthamiana* leaf extracts (peak area, transition *m/z* 331 > 216) transiently produced by co-infiltration of 4DO pathway genes. Empty vector (EV) was used as a negative control. The data was based on at least 3 biological replicates. Statistical analysis was performed using one-way ANOVA, where ∗ denotes significant difference ($p$<0.05), ∗∗ denotes extremely significant difference ($p$<0.01), and ns denotes no significant difference.

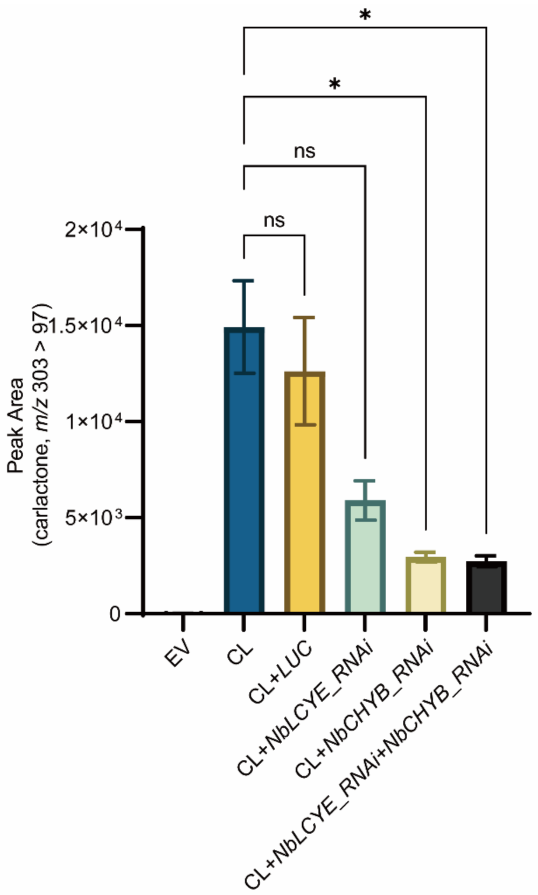


**figure S3. Effect of transient silencing of endogenous competing pathways of β-carotene in *Nicotiana benthamiana* on carlactone production**. Quantification of carlactone (CL) in *N. benthamiana* leaf extracts (peak area, transition *m/z* 303 > 97) transiently produced by co-infiltration of CL pathway genes and the silencing constructs *RNAi_NbLCYE* and *RNAi_NbCHYB1*. *RNAi_LUC* is a control silencing construct with a fragment of the firefly luciferase gene (Cankar et al., 2015). Empty vector (EV) was used as a negative control. The data was based on at least 3 biological replicates. Statistical analysis was performed using one-way ANOVA, where ∗ denotes significant difference ($p<0.05$), ∗∗ denotes extremely significant difference ($p<0.01$), and ns denotes no significant difference.
